# Single-Cell Multi-Omics Insights into BMI-Mediated Immune-Related Disease Risk

**DOI:** 10.64898/2026.01.12.698924

**Authors:** Zhuoli Huang, Zekai Xu, Han Yang, Yuhui Zheng, Wenwen Zhou, Baibing Guan, Yin Zeng, Jiaqiang Zhang, Pengbin Yin, Chuanyu Liu, Jianhua Yin

**Affiliations:** State Key Laboratory of Genome and Multi-omics Technologies, BGI Research, Shenzhen, China; College of Life Sciences, University of Chinese Academy of Sciences, Beijing, China; School of Biology and Biological Engineering, South China University of Technology, Guangzhou, China; School of Life Sciences, Southwest University, Chongqing, China; Department of Anesthesiology and Perioperative Medicine, People’s Hospital of Zhengzhou University, Henan Provincial People’s Hospital, People’s Hospital of Henan Academy of Innovations in Medical Science, Zhengzhou, China; Institute of Electrophysiology, Henan Academy of Innovations in Medical Science, Zhengzhou, China; Department of Orthopedics, Chinese PLA General Hospital, Beijing, China; National Clinical Research Center for Orthopedics, Sports Medicine & Rehabilitation, Beijing, China; Shenzhen Proof-of-Concept Center of Digital Cytopathology, BGI Research, Shenzhen, China; Shanxi Medical University-BGI Collaborative Center for Future Medicine, Shanxi Medical University, Taiyuan, China; Zhangzhou Affiliated Hospital of Fujian Medical University, Zhangzhou, China

## Abstract

This study is based on the Chinese Immune Multi-Omics Atlas (CIMA) cohort and performs an integrated single-cell multi-omics analysis of peripheral blood mononuclear cells (PBMCs) from 210 individuals aged 20-29, comprising 3,311,699 scRNA-seq cells and 1,839,860 scATAC-seq cells, to decipher immune characteristics across different body mass index (BMI) categories. By integrating gene regulatory network (GRN) and cell type-level molecular quantitative trait loci (xQTL) data, and conducting summary-data-based Mendelian randomization (SMR) analysis in conjunction with genome-wide association studies (GWAS) of BMI-mediated immune-related diseases, we identified key disease-associated cell types and molecular features. Using rheumatoid arthritis (RA) as an example, multi-omics evidence revealed that in MAIT-SLC4A10 cells of underweight individuals, increased chromatin accessibility at the *CCR6* locus (chr6:167119627-167120128) and enhanced transcriptional activity of RORC/RORA collectively drive elevated *CCR6* expression, which may represent a potential molecular mechanism underlying the higher risk of RA in underweight populations. Our study leverages single-cell multi-omics data to systematically dissect the molecular mechanisms by which different BMI categories mediate susceptibility to immune-related diseases through coordinated regulation of immune cell chromatin accessibility, transcription factor activity, and downstream gene expression.

## INTRODUCTION

The Body Mass Index (BMI) is a widely used metric that calculates an individual’s body weight relative to their height, commonly employed to classify adults into different body mass categories ^1^. While typically seen as an index of fatness, BMI is also a key risk factor for the development and prevalence of various health issues, influencing both individual health assessments and public health policies. Its widespread acceptance makes it an essential tool in population-based studies on body mass and health ^2,3^.

Variation in BMI is highly heritable and reflects a strongly polygenic architecture ^4,5^. To date, large-scale genome-wide association studies (GWAS) have systematically characterized the genetic basis of BMI ^6–9^. For instance, a comprehensive analysis involving approximately 700,000 individuals revealed 941 BMI-associated SNPs, which collectively explained about 6.0% of the BMI variance ^7^. Another study, based on data from 718,734 individuals and identified 14 rare and low-frequency coding variants in 13 genes associated with BMI ^8^. These findings provide a solid foundation for understanding how BMI may contribute to disease risk. Leveraging BMI-associated genetic variants as instrumental variables, numerous Mendelian randomization (MR) studies have been conducted to examine the potential causal effects of BMI on a broad range of complex diseases, including immune-related disorders ^4,10–17^. A representative study ^4^ demonstrated, through a phenome-wide MR analysis, that genetically predicted BMI was associated with 316 phecodes across 16 distinct disease categories. However, these studies have not provided insights into how different BMI categories affect disease risk at the molecular level. While GWAS and MR studies highlight statistical associations between BMI and various disease outcomes, they fall short of elucidating the cellular and molecular mechanisms that drive these associations.

Single-cell multi-omics technologies, including single-cell RNA sequencing (scRNA-seq), chromatin accessibility (scATAC-seq), and immune receptor profiling, are advancing the study of diseases by providing high-resolution insights into cellular heterogeneity and molecular mechanisms ^18–20^. These technologies enable the dissection of important cell types and their intricate regulatory networks, revealing how distinct cellular states contribute to disease pathogenesis ^20–22^. For example, in sepsis research, single-cell multi-omics profiling of peripheral blood mononuclear cells (PBMCs) identified specific immune cell subsets, such as NR4A2^+^ central memory CD4^+^ T cells, that are enriched in distinct sepsis sites and linked to immune exhaustion ^21^. Another study integrates GWAS with scRNA-seq and scATAC-seq data, found that fetal photoreceptor cells to be associated with major depressive disorder, and adult VGLUT2 neurons with schizophrenia, demonstrating how such integrative approaches can deepen our understanding of disease mechanisms ^22^. Currently, single-cell studies on BMI primarily focus on adipose tissue ^23–25^, with limited attention given to the differences in immune cell types and characteristics in PBMCs across different BMI categories. Additionally, both underweight and obesity are associated with adverse health outcomes throughout the life course ^26^, yet many studies on BMI tend to focus predominantly on obesity or overweight, often neglecting the implications of being underweight.

To address these issues, our study builds upon the previously published Chinese Immune Multi-Omics Atlas (CIMA) cohort ^27^. We selected 210 samples from individuals aged 20-29, comprising 3,311,699 scRNA-seq cells and 1,839,860 scATAC-seq cells, and utilized the gene regulatory network (GRN) from this study to perform an integrated multi-omics analysis of immune characteristics across different BMI categories. Additionally, leveraging the cell type-level molecular quantitative trait loci (xQTL) data from this cohort, we conducted a joint analysis with GWAS data of BMI-mediated immune-related diseases to identify disease-associated cell types and molecular features. Building on the findings from these analyses, we aim to elucidate how distinct BMI categories influence the downstream risk of immune-related diseases through modulation of chromatin accessibility and gene expression in immune cells. This research provides valuable insights into the molecular mechanisms through which BMI affects immune-related diseases.

## RESULTS

### Overview of the Study Cohort

The study cohort used in this research is part of the CIMA cohort ^27^. Specifically, the CIMA provides scRNA-seq and scATAC-seq data across 73 cell types from 428 donors, along with GRNs and cell type-level xQTL data. From the scRNA-seq and scATAC-seq datasets, we selected 210 donors aged 20–29 years (Supplementary Table 1), accounting for 49.06% of the total samples, which comprised 3,311,699 scRNA-seq cells and 1,839,860 scATAC-seq cells. These data were integrated with GRN and xQTL resources for comprehensive analysis (Methods, Fig. 1a).

**Figure. 1.**
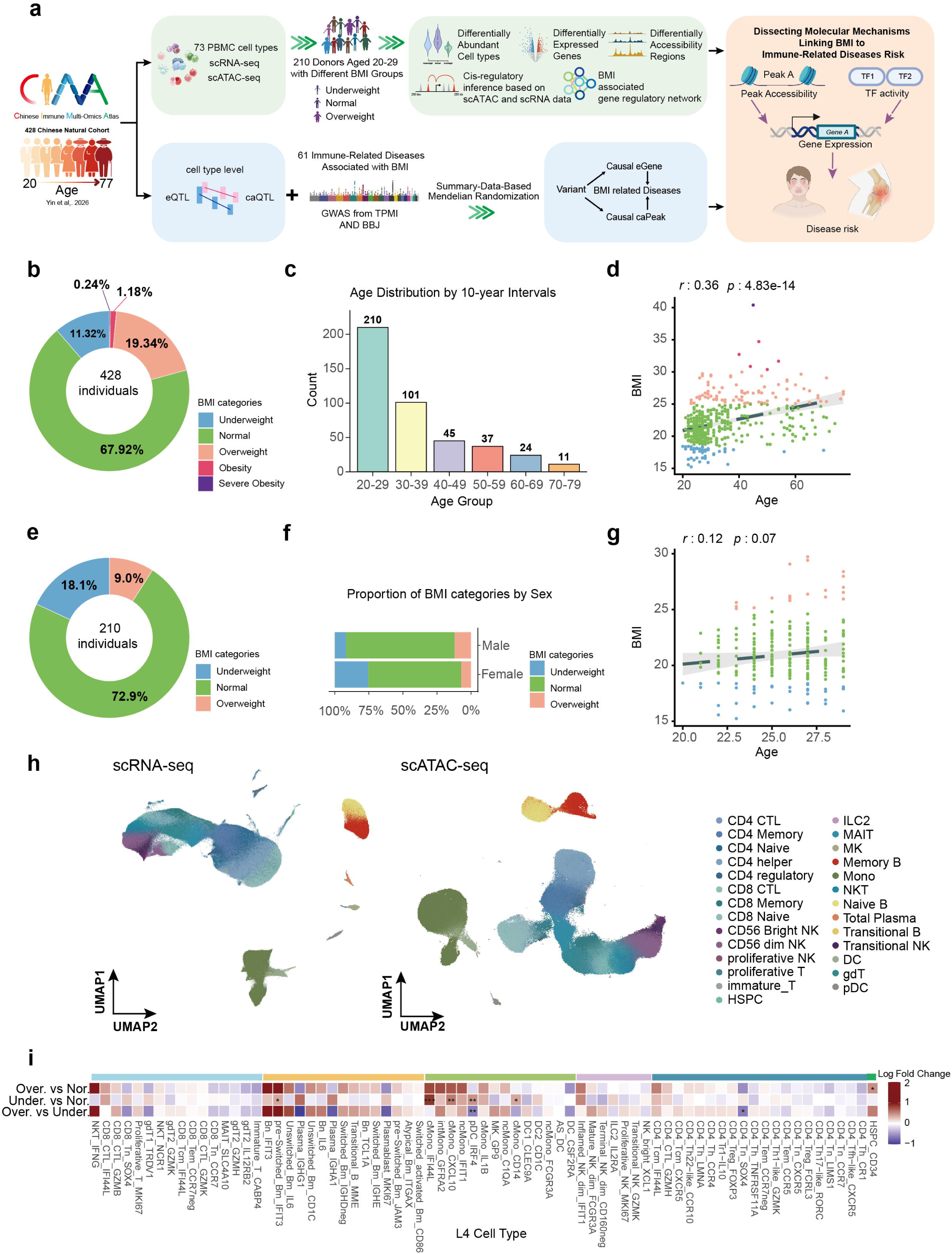
Study design and overview of cohort categorized by BMI categories. (a) Overview of the selected CIMA cohort and analysis workflow. (b) The pie chart shows the distribution of BMI categories in the CIMA cohort. (c) Bar plot illustrating the number of individuals in the CIMA cohort across different age Intervals. (d) Linear regression analysis estimating the relationship between age and BMI in the CIMA cohort. Each dot represents one individual. (e) The pie chart shows the distribution of BMI categories in the selected CIMA cohort aged 20-29 years. (f) Stacked bar plot illustrating proportion of BMI categories by sex. (g) Linear regression analysis estimating the relationship between age and BMI in the selected CIMA cohort aged 20-29 years. Each dot represents one individual. (h) UMAP plot showing immune cells from the scRNA-seq data and scATAC-seq in the selected CIMA cohort aged 20-29 years, based on Level 2 annotation of cell types. (i) Heatmap shows differences in cell type proportions across BMI categories.

Participants in the CIMA cohort all self-reported no active disease at sampling ^27^, resulting in a very low prevalence of obesity (1.18%) and severe obesity (0.24%) (Fig. 1b). The cohort age range is 20-77 years, with the highest proportion of individuals aged 20-29 (Fig. 1c). Previous research has demonstrated an inverted U-shaped trajectory of BMI across the lifespan ^28^, and a similar age-related trend was observed in the CIMA cohort, where BMI was positively correlated with age in a linear regression analysis (Fig. 1d). Since age significantly confounds immune state ^29,30^, analyzing BMI categories across the full cohort would conflate age-related effects. Therefore, to specifically isolate BMI-associated effects, we focused on the 20-29 age group. Within these 210 individuals, the underweight, normal, and overweight categories accounted for 18.1%, 72.9%, and 9.0% of the subset, respectively (Fig. 1e). All three BMI categories were represented in both males and females (Fig. 1f), with no significant association observed between BMI and age in this cohort (Fig. 1g). For cell type annotation, we adopted the hierarchical annotation framework comprising four levels from the CIMA cohort ^27^. This framework encompasses 6 Level 1 cell types, 27 Level 2 cell types (Fig. 1h), 38 Level 3 cell types, and 73 Level 4 (L4) cell types. For all downstream analyses, we used the most granular L4 annotations. Among the 73 cell types, the proportions of 7 cell types were detected to differ across BMI categories. Specifically, compared to the normal category, the underweight category showed significantly elevated proportions of pre-Switched-Bm-IFIT3, cMono-CD14, cMono-CXCL10, cMono-IFI44L, and pDC-IRF4. In contrast, the proportion of HSPC-CD34 was significantly increased in the overweight category (Fig. 1h).

### Single-cell multi-omics analysis of the PBMC in different BMI categories

To investigated BMI-associated gene expression and regulatory chromatin by cell type, we analyzed scRNA-seq and scATAC-seq from the PBMC of 210 donors. Differential gene expression analysis was performed across all 73 cell types using the Wilcoxon rank-sum test (Methods). Differentially expressed genes (DEGs) were identified in 72 of these cell types (Supplementary Table 2). In the comparison between the overweight and normal categories, we identified 2,806 DEGs across 68 cell types, of which 35.9% (1,035 genes) were differentially expressed in only a single cell type (Fig. 2a). In the comparison between the underweight and normal categories, we detected 1,143 DEGs across 71 cell types, with 63.5% (726 genes) showing differential expression in just one cell type (Fig. 2b). When comparing underweight to overweight category, we found 3,351 DEGs across 70 cell types, and 40.5% (1,358 genes) of these were uniquely dysregulated in a cell type (Supplementary Fig. 1a). Notably, Mature-NK-dim-FCGR3A, cMono-CD14, gdT2-GZMH, and CD4-Tn-CCR7 exhibited relatively higher numbers of DEGs.

**Figure. 2.**
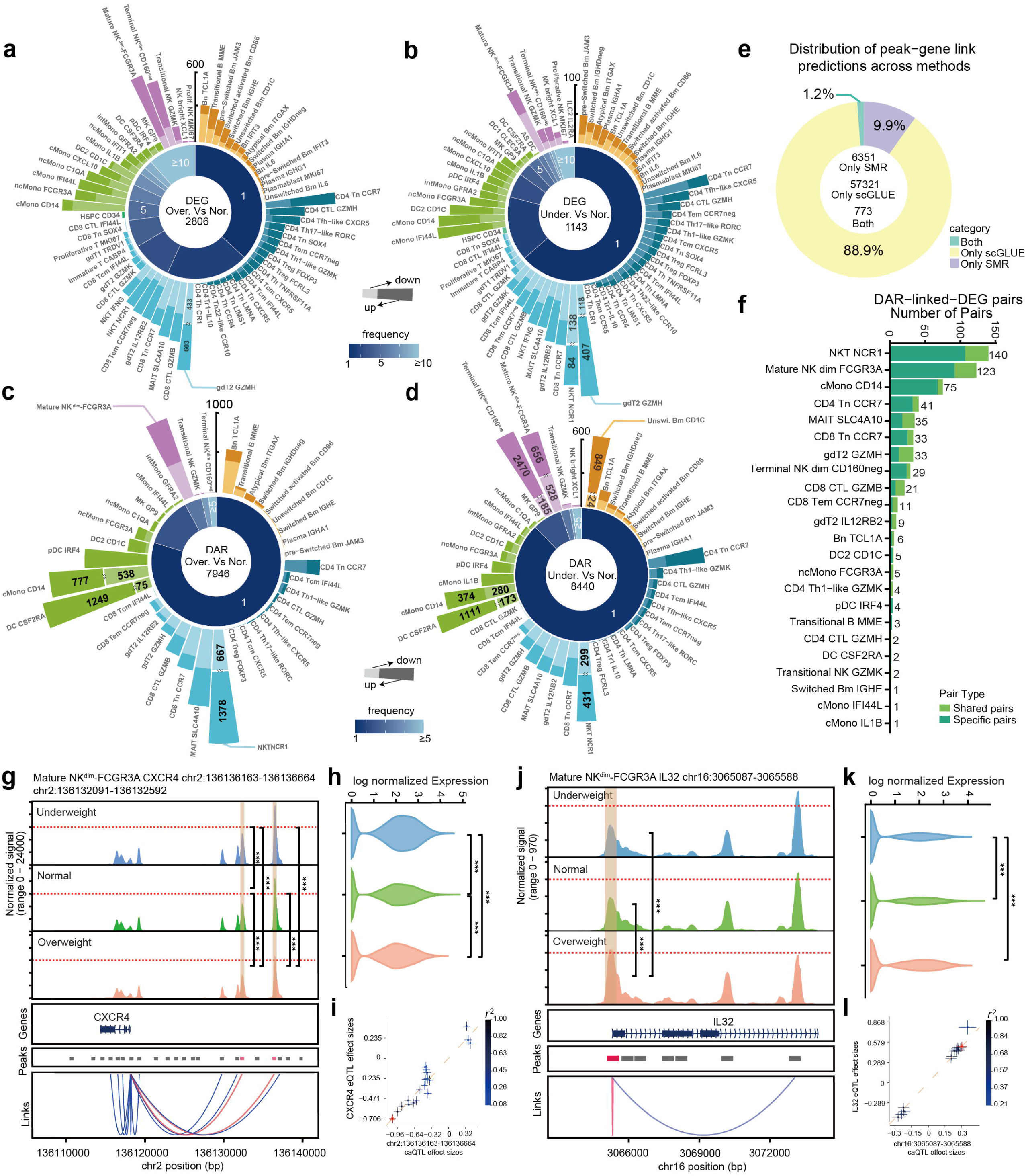
Epigenetic dysregulation in BMI-related peripheral immunity and concordance with differential gene expression. (a and b) Bar plot illustrating the number of DEGs detected across 68 cell types (a) and 71 cell types (b) in different BMI categories. The pie chart shows the distribution of significantly detected DEGs, categorized by the number of cell types in which each DEG was identified. (c and d) Bar plot illustrating the number of DARs detected across 38 cell types (c) and 44 cell types (d) in different BMI categories. The pie chart shows the distribution of significantly detected DARs, categorized by the number of cell types in which each DAR was identified. (e) The pie chart displays the distribution of significantly detected peak-gene links. (f) Bar plot illustrating the number of significant peak-gene links which peak is DAR and gene is DEG across the 23 cell types. (g) Chromatin track of chr2:136136163-136136664 and chr2:136132091-136132592 intersecting *CXCR4* gene in different BMI categories in Mature NKdim-FCGR3A. The DARs of interest are highlighted in orange. The linkage plot illustrates associations with chromatin accessibility and gene expression which were identified through either SMR-based colocalization or cis-regulatory inference using scGLUE feature embeddings. The link of interest is highlighted in red. (h) Violin plot shows *CXCR4* expression in different BMI categories in Mature NKdim-FCGR3A. (i) Effect sizes of variants from chr2:136136163-136136664 caQTL plotted against *CXCR4* eQTL variants in Mature NKdim-FCGR3A. (j) Chromatin track of chr16:3065087-3065588 intersecting *IL32* gene in different BMI categories in Mature NKdim-FCGR3A. The DARs of interest are highlighted in orange. The linkage plot illustrates associations with chromatin accessibility and gene expression which were identified through either SMR-based colocalization or cis-regulatory inference using scGLUE feature embeddings. The link of interest is highlighted in red. (k) Violin plot shows *IL32* expression in different BMI categories in Mature NKdim-FCGR3A. (l) Effect sizes of variants from chr16:3065087-3065588 caQTL plotted against *IL32* eQTL variants in Mature NKdim-FCGR3A.

Circulating classical monocytes (cMono) are closely associated with BMI ^31,32^. We therefore performed Gene Ontology (GO) enrichment analysis on DEGs in cMono-CD14 (Method and Supplementary Table 3). Compared to the normal category, the overweight category exhibited 87 upregulated genes, which were significantly enriched in pathways related to type II interferon response, tumor necrosis factor (TNF) production, and TNF superfamily cytokine production (Supplementary Fig. 1c). Meanwhile, 88 downregulated genes were enriched in MHC class II-mediated antigen presentation pathways (Supplementary Fig. 1d). In contrast, compared to the normal category, the underweight category showed 70 upregulated genes, enriched in immune response-regulating cell surface receptor signaling pathways, as well as antigen recognition and binding pathways (Supplementary Fig. 1e). Only 42 genes were downregulated, and due to the limited number, no significant functional enrichment was observed.

To identify differentially accessible peaks across BMI categories, differential accessibility analysis was performed across all 64 cell types using the logistic regression (LR) method (Methods). Differentially accessible regions (DARs) were detected in 44 cell types (Supplementary Table 4). In the comparison between the overweight and normal categories, we identified 7,946 DARs across 38 cell types, of which 79.7% (6,333 peaks) were differentially accessible in only one cell type (Fig. 2c). In the underweight versus normal comparison, we detected 8,440 DARs across 44 cell types, with 87.7% (7,404 peaks) unique to one cell type (Fig. 2d). Comparing underweight to overweight category, we found 7,500 DARs across 39 cell types, and 77.4% (5,802 peaks) were specific to one cell type (Supplementary Fig. 1b). Notably, Mature-NK-dim-FCGR3A, cMono-CD14, gdT2-GZMH, and CD4-Tn-CCR7 exhibited relatively higher number of DARs.

We further investigated functional differences at the epigenomic level. Associating each DAR with its location within the genome revealed that most DARs were located in promoter and intronic regions (Supplementary Fig. 2a). We next focused on cMono-CD14 and performed GO enrichment analysis of DARs using rGREAT (Methods and Supplementary Table 5). Compared to the normal category, the overweight category exhibited 777 more accessible peaks, which were enriched in kinase-related pathways and interleukin-1 receptor binding. Meanwhile, 538 less accessible peaks were enriched in various signal transduction-related pathways (Supplementary Fig. 2b). Compared to the normal category, the underweight category showed 374 more accessible peaks, enriched in kinase signaling pathways and signal receptor transduction-related processes. A total of 280 less accessible peaks were significantly enriched in chemokine-related pathways (Supplementary Fig. 2c).

To further identify reliable and interpretable multicellular pathway modules and capture molecular feature aberrations across individuals with different BMI categories, we applied single cell pathway activity factor analysis (scPAFA) ^33^ (Methods). We performed scPAFA ^33^ using gene expression data across immune cell types, identifying factor 4 as the most BMI-associated latent component. The value of Factor 4 was significantly higher in overweight than other two categories, while no significant differences were observed between sexes (Supplementary Fig. 3a, b). We further employed the feature weights matrixes corresponding to factors 4 to interpret BMI-related multicellular pathway modules (Supplementary Fig. 3c). We observed that pathways with high positive loadings on factor 4 were enriched for type I interferon response across multiple cell types. This pattern was consistent with the functional enrichment results of DARs and suggested a more active metaflammatory state in overweight individuals. In contrast, pathways with high negative loadings on factor 4 included mitophagy regulation–related pathways in CD4-Tn-CCR7, CD8-Tn-CCR7, and Bn-TCL1A, as well as lactate production pathways in CD8-Tn-CCR7 and Bn-TCL1A. Mitochondrial dysfunction is recognized as a key contributor to the pathogenesis of obesity and other metabolic disorders ^34^. To maintain metabolic homeostasis, mitochondrial quality is tightly regulated through the coordinated actions of selective mitochondrial autophagy, mitochondrial biogenesis, and multiple mitochondrial protein degradation processes ^35,36^. The relationship between lactate and BMI is complex. Previous studies have suggested that lactate may act as a critical mediator linking obesity to insulin resistance. However, the association between lactate metabolism in PBMCs and BMI remains unexplored ^37^. Together, our findings may reflect BMI-associated alterations in energy metabolism within PBMC.

### Cis-regulation of gene expression associated with BMI

To interrogate the relationship between BMI and the peripheral immune system, we integrated scGLUE ^38^-based cis-regulatory inferences on the data and summary-data-based Mendelian randomization (SMR) ^39^ analyses of caQTLs and eQTLs from CIMA to compile associations between chromatin peaks and their neighboring genes (referred to as peak-gene links) (Methods). Finally, we identified 64,445 peak-gene links, of which 88.9% were significant only in the scGLUE ^38^ analysis, 9.9% were significant only in the SMR analysis, and 1.2% were significant in both approaches (Fig. 2e; Supplementary Table 6). By integrating the identified peak-gene links with our previously obtained BMI-associated DARs and DEGs, we established regulatory relationships between BMI-associated peaks and their target genes. This integration yielded a total of 586 DAR-linked-DEG pairs across 23 cell types, with NKT-NCR1 containing 140 such pairs and Mature-NK-dim-FCGR3A containing 123 pairs (Fig. 2f).

Notably, we identified some BMI-associated peak-to-gene links that are shared across multiple cell types. An example is the observation of BMI-associated regulation of the C-X-C motif chemokine receptor 4 (CXCR4) gene in Mature-NK-dim-FCGR3A and NKT-NCR1. Compared with normal, *CXCR4* was upregulated in underweight (log_2_FC=0.31, *p*_adj_=9.23e-127) and was downregulated in overweight (log_2_FC=-0.36, *p*_adj_ =2.62e-104) in Mature NK dim-FCGR3A. We observed 2 promoter regions chr2:136136163-136136664 and chr2:136132091-136132592 probably influencing the expression of *CXCR4*, and chromatin accessibility at these two regions exhibited significant differences across BMI categories (Fig. 2g, h). The chr2:136136163-136136664 and chr2:136132091-136132592, showed significant associations with *CXCR4* gene expression, supported by the eQTL and caQTL analyses using the SMR pleiotropy test from CIMA (Figure 2i, and Supplementary Table 6). The SMR pleiotropy test revealed a positive association between chromatin accessibility at the two peaks and *CXCR4* expression levels, corroborating our result. A similar pattern was also observed in NKT-NCR1 (Supplementary Fig. 4a).

CXCR4 plays a key role in NK cell trafficking and function ^40^, and in adipose tissue, the CXCL12-CXCR4 signaling pathway has been linked to insulin resistance ^41^. Overall, these findings suggest a shared BMI-associated immunoregulatory mechanism in Mature-NK-dim-FCGR3A and NKT-NCR1.

Another example is the BMI-associated regulation of the Interleukin 32 (IL32) gene in the Mature-NK-dim-FCGR3A. IL32 expression was significantly downregulated in the underweight group compared to both the normal (log2FC = -0.26, *p*_adj_ = 1.15e-35) and overweight groups (log2FC = -0.44, *p*_adj_ = 2.73e-52) (Fig. 2k). We observed a promoter regions chr16:3065087-3065588 probably influencing gene expression, and chromatin accessibility at these two regions exhibited significant differences across BMI categories (Fig. 2j, k). The SMR pleiotropy test form CIMA revealed a positive association between chromatin accessibility at the chr16:3065087-3065588 and *IL32* expression levels, corroborating our result (Fig. 2l, and Supplementary Table 6). Previous studies have demonstrated that IL-32 is involved in the pathophysiological disturbances within adipose tissue ^42^, and *IL32* expression is upregulated in PBMC of individuals with obesity ^43^, suggesting a role for *IL32* in BMI-associated inflammatory regulation. Our findings further indicate that *IL32* in Mature NK dim-FCGR3A may contribute to this inflammatory response, potentially mediated by the region of chr16:3065087-3065588.

*HLA-DQB1* exhibits a BMI-associated regulatory pattern similar to the examples described above in gdT2 GZMH. However, no significant SMR association was observed. Only a cis-regulatory link inferred by scGLUE ^38^ supports this relationship (Supplementary Fig. 4b). A previous study reported that *HLA-DQB1* expression in PBMC is downregulated in individuals with obesity compared to those who are underweight ^44^. HLA gene family is well known for its central role in antigen presentation and immune regulation, the specific mechanisms underlying its interaction with BMI remain poorly characterized. In summary, we have identified a series of potential BMI-associated gene regulatory relationships in PBMC.

### BMI-associated GRN

Transcription factors (TFs) activity is an important molecular feature, as TFs influence chromatin accessibility and mediate gene expression ^45^. To elucidate the relationship between BMI-associated TFs and downstream transcriptional alterations, we conducted TF enrichment analyses of BMI-associated DEGs and DARs using 203 GRN-defined enhancer-linked regulatory units (eRegulons) from the CIMA cohort, each representing a TF-peak-gene regulatory module. This analysis identified 97 TFs that were significantly enriched (Methods; Fig. 3a, b; Supplementary Table 7 and 8). Among the significantly enriched factors were several immune regulatory TFs, including FOSB, JUNB, JUN, and XBP1.

**Figure. 3.**
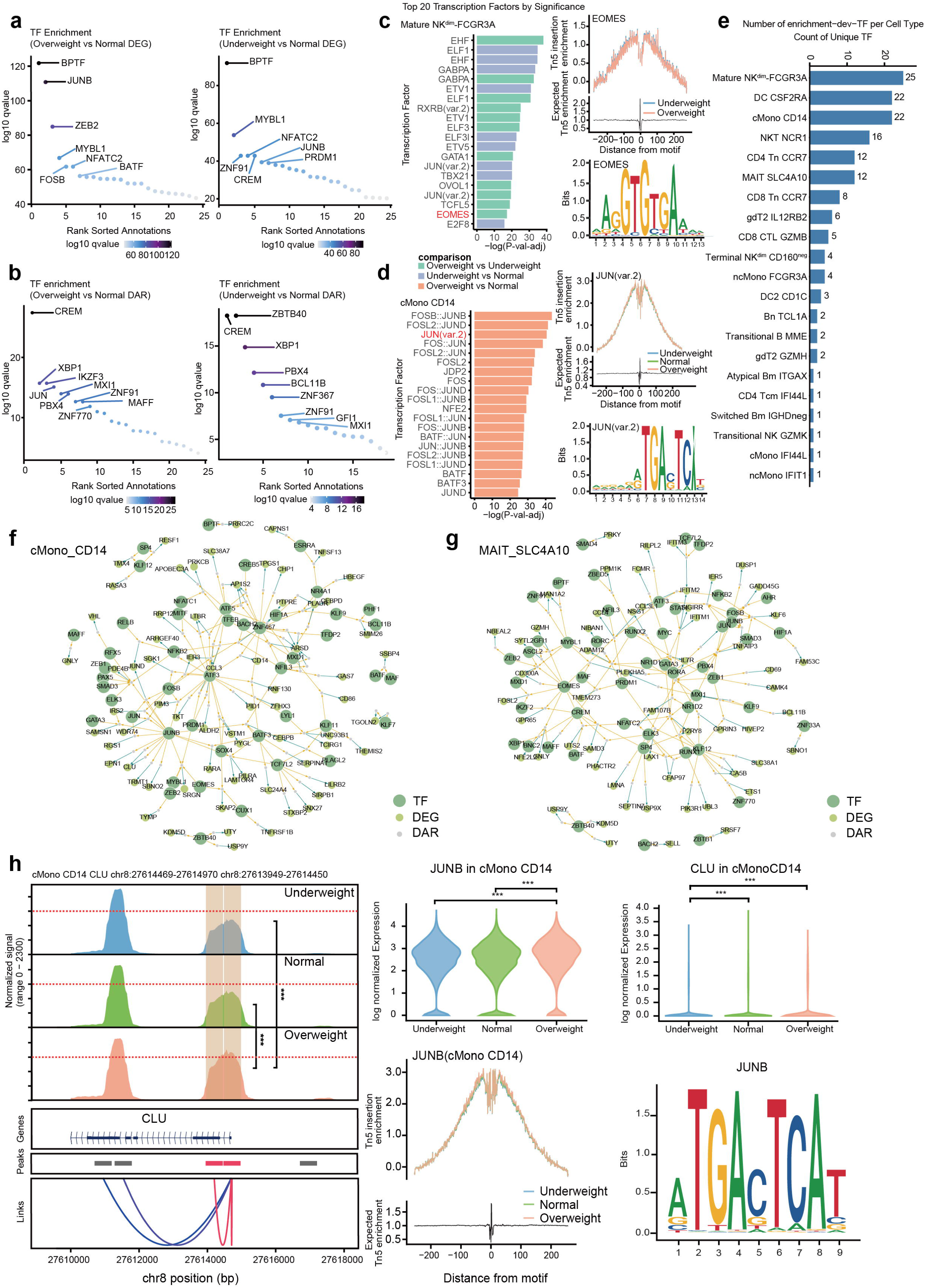
The role of TFs in BMI. (a) TF enrichment in DEGs (Overweight vs Normal (left) and Underweight vs Normal (right)). TFs labeled in the plot are those with the smallest adjusted enrichment p-values. (b) TF enrichment in DARs (Overweight vs Normal (left) and Underweight vs Normal (right)). TFs labeled in the plot are those with the smallest adjusted enrichment p-values. (c) Bar plot (left) showing the top TF motifs with the −log_10_(P) values, whose motif activity differs between BMI categories in Mature NKdim-FCGR3A. Tn5 bias-subtracted footprints of ELF1 in Mature NKdim-FCGR3A and DNA sequence logo of the ELF1 motifs (right). (d) Bar plot (left) showing the top TF motifs with the −log_10_(P) values, whose motif activity differs between BMI categories in cMono-CD14. Tn5 bias-subtracted footprints of JUN(var.2) in cMono-CD14 and DNA sequence logo of the JUN(var.2) motifs (right). (e) Bar plot showing the number of TFs that exhibit differential motif activity and are also enriched in TF enrichment analysis across 21 cell types. (f and g) Gene regulatory networks of BMI-associated prioritized TFs in cMono-CD14 (f) and MAIT-SLC4A10 (g). Based on SCENIC+-derived eRegulons, which define TF–peak–gene regulatory axes, the overlap between eRegulons containing significantly enriched TFs and cell-type–specific DEGs and DARs is shown. (h) Chromatin track of chr8:27614469-27614970 and chr8:27613949-27614450 intersecting CLU gene in different BMI categories in cMono-CD14. The DARs of interest are highlighted in orange (left). The linkage plot illustrates associations with chromatin accessibility and gene expression which were identified through either SMR-based colocalization or cis-regulatory inference using scGLUE feature embeddings. The link of interest is highlighted in red. Violin plot shows JUNB expression and CLU expression in different BMI categories in cMono-CD14 (right). Tn5 bias-subtracted footprints of JUNB in cMono-CD14 and DNA sequence logo of the JUNB motifs (right).

To further investigate BMI-associated differential TFs, we assessed TF motif deviation z-scores, corrected for GC content and chromatin accessibility biases, across BMI categories within each cell type using chromVAR (Methods; Fig. 3c, d). Overall, 590 TFs showed significant differences in binding activity among BMI categories (Supplementary Table 9). We observed that Mature-NK-dim-FCGR3A exhibited the largest number of TFs with significant differences in motif activity across BMI categories (Supplementary Fig. 5a). We focused on two cell types that showed strong associations with BMI comparisons: Mature-NK-dim-FCGR3A and cMono-CD14. In Mature-NK-dim-FCGR3A cells, the motif deviation z-score of ELF1 differed significantly among BMI categories (Fig. 3c), with higher predicted ELF1 binding activity in the underweight compared with the normal and overweight. ELF1 plays a critical role in the immune system and is involved in the transcriptional regulation of multiple T-cell---specific molecules, including IL2R ^46^. *ELF1* is also expressed in Mature-NK-dim-FCGR3A (Supplementary Fig. 5b), but previous studies have shown that ELF1 has minimal impact on NK cell development ^47^. Its potential role in modulating NK cell function across different BMI states warrants further investigation. In cMono-CD14, we observed higher predicted binding activity of JUN (var2) in the overweight category compared with the normal category (Fig. 3d). *JUN* is expressed in cMono-CD14 (Supplementary Fig. 5c) and forms heterodimers with members of the FOS family, thereby participating in the regulation of immune-related gene expression ^48^. By overlapping the results from chromVAR-based motif activity analysis and TF enrichment analysis, we identified TFs most likely associated with BMI. Among these, Mature-NK-dim-FCGR3A, cMono-CD14, and MAIT-SLC4A10 exhibited a relatively high number of BMI-associated TFs involved in gene regulation (Fig. 3e).

Furthermore, by intersecting cell type-specific BMI-associated DEGs and DARs with the eRegulons of the 97 significantly enriched TFs, we constructed BMI-associated GRN within immune cell type, with cMono-CD14 and MAIT-SLC4A10 shown as representative examples (Fig. 3f, g). We observed that, in cMono-CD14(Fig. 3f), several immune response-related TFs, including JUNB, JUN, and FOSB, exerted coordinated regulatory effects on subsets of DARs and DEGs. In MAIT-SLC4A10, TFs such as RORC and RORA, which are implicated in metabolic regulation, immune function, and circadian rhythm, were also associated with altered expression of target genes (Fig. 3g). One example in cMono-CD14 is that chromatin accessibility at JUNB TF binding sites differed across BMI categories and was associated with expression of the *CLU* gene (Fig. 3g).

Previous studies have shown that CLU protein can modulate insulin resistance and is associated with multiple cardiovascular disease and cardiovascular risk ^49,50^. We further observed that *JUNB* gene expression varied across BMI categories, accompanied by corresponding differences in predicted TF binding activity (Fig. 3h). Collectively, these results suggest a potential mechanism by which BMI-associated changes in TF activity and chromatin accessibility may influence to downstream gene expression.

### SMR Analysis Identifies Key Cell Types and Features Underlying BMI-Mediated Immune-Related Diseases

Based on the established cell type-level DAR and DEG GRN associated with BMI categories, we aim to further investigate cell types and immune signatures relevant to BMI-mediated immune-related diseases. Accordingly, leveraging GWAS data for 61 BMI-mediated immune-related diseases (Supplementary Table 10) in East Asian populations ^51,52^ and xQTL data from the CIMA cohort ^27^, we performed SMR analysis (Materials and Methods) to identify cell type-level pleiotropic associations of “variant–eGene/caPeak–disease.” The cell types and signatures detected through pleiotropic association analysis suggest potential causal links to the diseases.

Among the 68 cell types, we detected a total of 1,406 significant results (Supplementary Table 11), encompassing 864 pleiotropic associations, involving 84 eGenes, 146 caPeaks, and 25 traits (Fig. 4a). Of these 864 pleiotropic associations, 75.12% were specific to a single cell type, 23.73% were detected across 2 to 10 cell types, and 1.15% were observed in more than 10 cell types (Fig. 4b). Furthermore, among the 1,406 significant results, 8.96% overlapped with BMI categories-associated DEGs, and 1.56% overlapped with DARs within their corresponding cell types (Fig. 4b). The top three cell types ranked by number of pleiotropic associations were cMono-CD14, Mature-NK-dim-FCGR3A, and CD8-Tn-CCR7 (Fig. 4c); the top three diseases were Rheumatoid arthritis (RA), Asthma, and Psoriasis and related disorders (Fig. 4d). Existing study ^53^ on Chinese populations have shown that underweight individuals have a higher risk of developing RA. Taking RA as an example, its associated pleiotropic links involved 19 eGenes, 7 of which overlapped with BMI-related DEGs in the relevant cell types (Fig. 4e). These findings suggest that different BMI categories may influence disease risk through cell type-specific differences in gene expression.

**Figure. 4.**
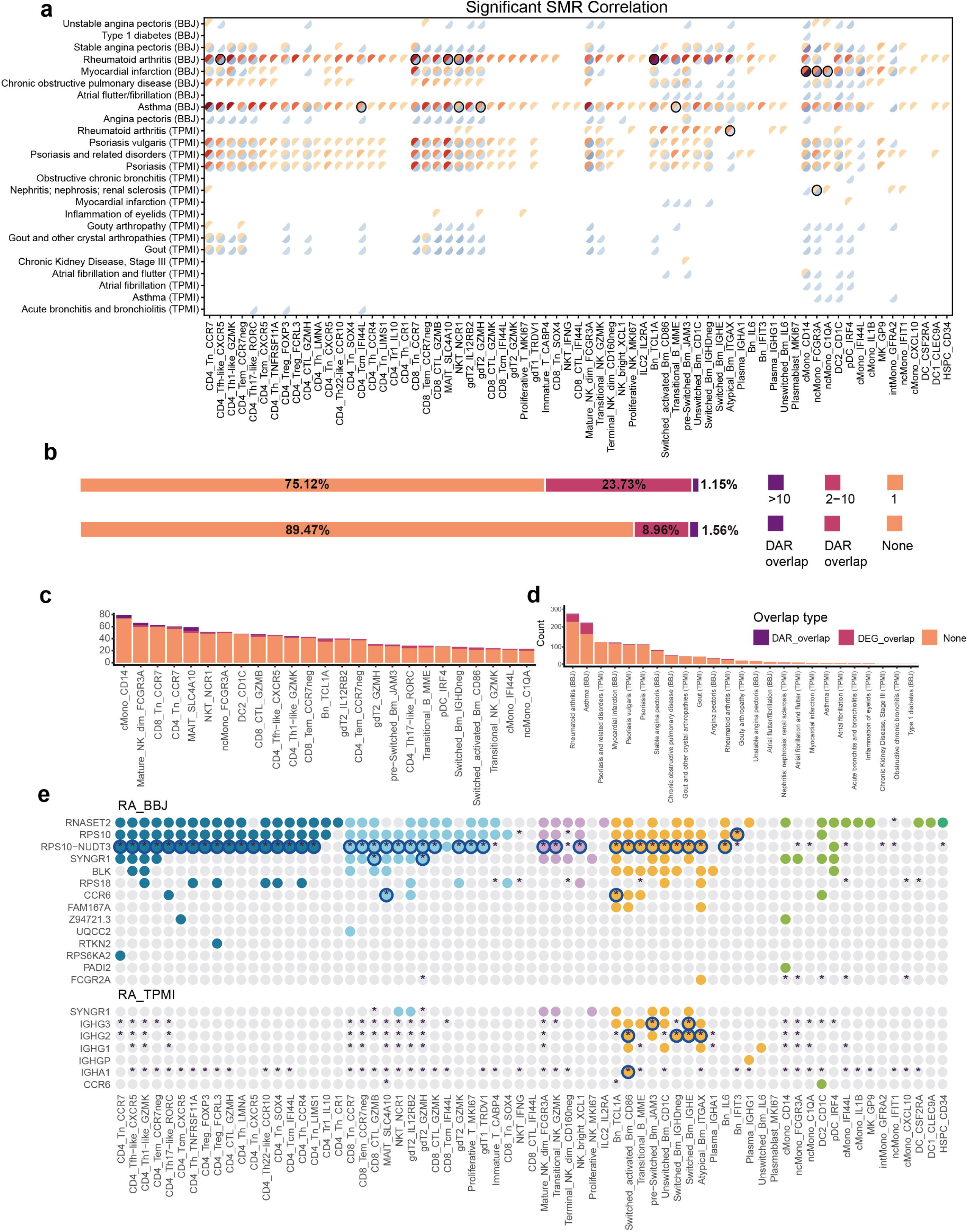
Summary of genetic pleiotropic associations. (a) Heatmap illustrating the number of eGenes and caPeaks detected in significant variant-eGene/caPeak-trait SMR pleiotropic associations across cell types. The black outline indicates that the shared top variant simultaneously pleiotropically associated with cis-eGene, cis-caPeaks and trait was detected. BBJ, Biobank Japan. TPMI, Taiwan Precision Medicine Initiative. (b) Stacked bar charts illustrating the distinct and shared patterns of significant SMR correlations among cell types (top), together with the overlap between SMR correlations and DEGs or DARs (bottom) (c) Bar plot showing the number of significant SMR correlations across 25 cell types for the top SMR signals, and their overlap with BMI-related DEGs and DARs. (d) Bar plot showing the number of significant SMR correlations across 25 BMI-related diseases, and their overlap with BMI-related DEGs and DARs. BBJ, Biobank Japan. TPMI, Taiwan Precision Medicine Initiative. (e) The eGenes detected in significant variant-eGene-disease SMR pleiotropic associations across cell types in Rheumatic Arthritis (RA) in BBJ and TPMI. Asterisks indicate that the gene is DEG in the corresponding cell type. The black outline indicates that the gene is DEG and shows pleiotropic association between rheumatoid arthritis (RA) GWAS and eQTL.

### Multi-Omics Dissection of Rheumatoid Arthritis Risk in the Underweight category

We further investigated using RA and *CCR6* as an example. *CCR6* showed pleiotropic associations with RA in both MAIT-SLC4A10 and Bn-TCL1A, with a lower *p*_eQTL_ in MAIT-SLC4A10 than in Bn-TCL1A, indicating a more significant effect (Fig. 5a). The effect size plot revealed a positive correlation between the eQTL effect size of *CCR6* and the GWAS effect size of RA, suggesting that *CCR6* expression may be positively associated with RA disease risk (Fig. 5b, c). In MAIT-SLC4A10, *CCR6* expression was significantly higher in the underweight category compared to the overweight category, whereas in Bn-TCL1A, *CCR6* expression in the overweight category was significantly lower than in the normal category. Since *CCR6* expression was higher in MAIT-SLC4A10 than in Bn-TCL1A, we focused on MAIT-SLC4A10. Further analysis showed that the high expression of *CCR6* in the underweight category within MAIT-SLC4A10 cells is supported by multi-omics evidence. In MAIT-SLC4A10 cells, the chr6:167119627-167120128 peak is linked to *CCR6* through both peak-to-gene regulatory relationships and SMR pleiotropic associations (Fig. 5e, f), with a positive correlation between caQTL and eQTL effects (Fig. 5e), suggesting that higher chromatin accessibility may enhance gene expression. As expected, the chr6:167119627-167120128 peak in MAIT-SLC4A10 cells showed significantly higher accessibility in the underweight category compared to the overweight category (Fig. 5f). Supporting this finding, the chr6:167119627-167120128 peak is located within a promoter/enhancer region in an intron of *CCR6* (Supplementary Fig. 6a).

**Figure. 5.**
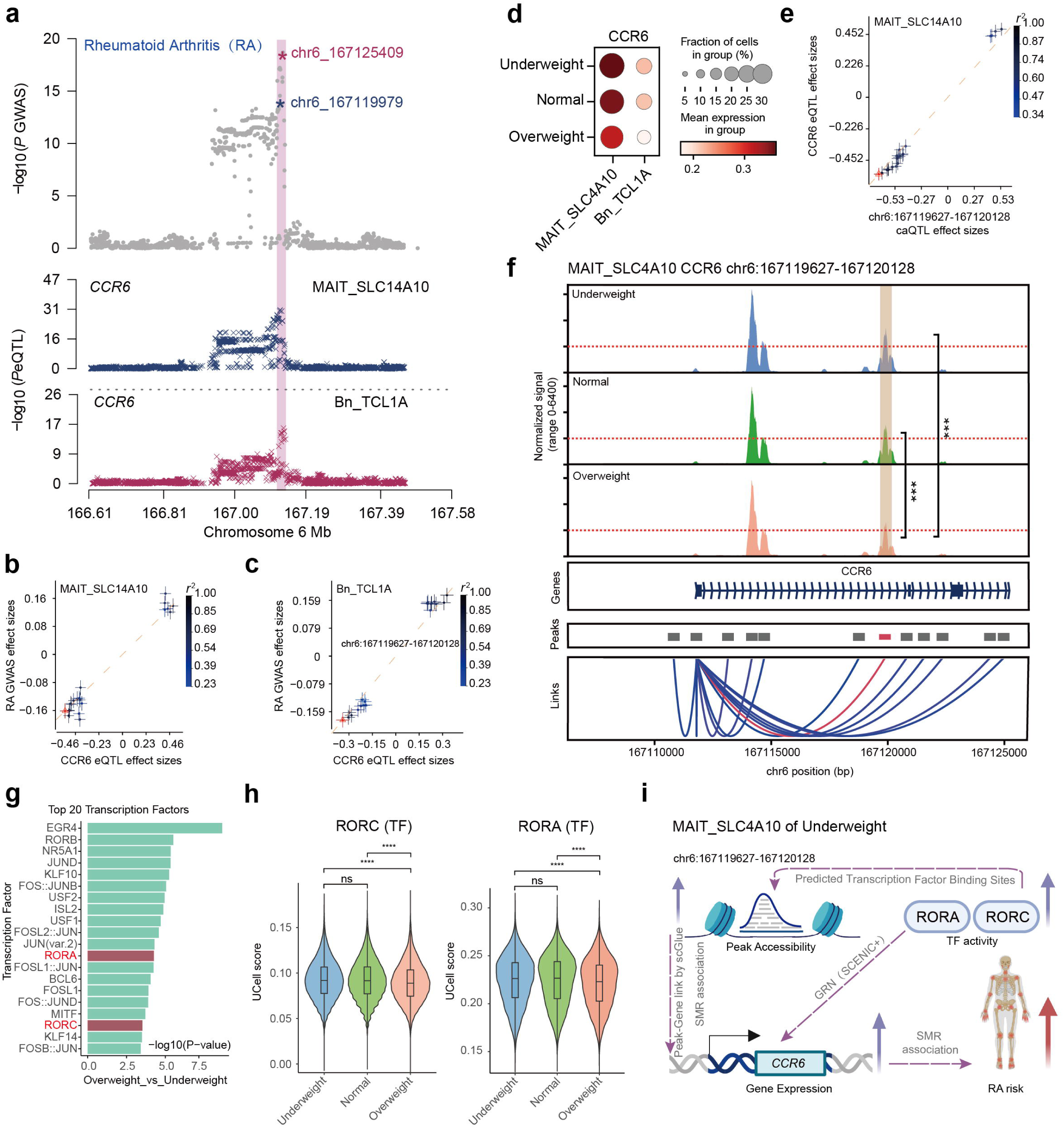
BMI-related cell-shared genetic pleiotropic associations. (a) Locus plots of significant SMR correlations among *CCR6* and RA in two relevant cell types in MAIT-SLC4A10 and Bn-TCL1A. (b and c) Effect sizes of variants from RA GWAS plotted against *CCR6* eQTL variants in MAIT-SLC4A10 (b) and in Bn-TCL1A (c). (d) Dot plot of *CCR6* expression across cell types. (e) Effect sizes of variants from chr6:167119627-167120128 caQTL plotted against *CCR6* eQTL variants in MAIT-SLC4A10. (f) Chromatin track of chr6:167119627-167120128 intersecting CCR6 gene in different BMI categories in MAIT-SLC4A10. The DARs of interest are highlighted in orange. The linkage plot illustrates associations with chromatin accessibility and gene expression which were identified through either SMR-based colocalization or cis-regulatory inference using scGLUE feature embeddings. (g) Bar plot showing the top TF motifs with the −log_10_(P) values, whose motif activity differs between Overweight and Underweight in MAIT SLC4A10. (h) Violin plot showing UCell score of RORC (left) and RORA (right) across different BMI categories in MAIT-SLC4A10. (i) Proposed mechanistic model depicting the effect of BMI-related *CCR6* expression to RA disease risk in MAIT-SLC4A10.

Moreover, the BMI-related GRNs of MAIT-SLC4A10 includes the TFs RORC and RORA (Fig. 3g), which are known upstream regulators of *CCR6* ^54–56^, where increased transcriptional activity can elevate *CCR6* expression. TF deviation analysis indicated significant differences in RORC and RORA activity between the underweight and overweight category (Fig. 5g). Based on SCENIC+ ^57^ TF Ucell scores, the transcriptional activities of RORC and RORA were significantly higher in the underweight category than in the overweight category (Fig. 5h). Additionally, JASPAR ^58^ predicted TF binding sites for RORC and RORA were identified within the chr6:167119627-167120128 peak region (Supplementary Fig. 6b). In summary, in the underweight category, higher chromatin accessibility at the chr6:167119627-167120128 peak and increased transcriptional activity of RORC and RORA may collectively enhance *CCR6* expression, potentially leading to higher RA disease risk (Fig. 5i). This finding further supports and explains the higher RA risk observed in underweight individuals in the Chinese population, building upon previous research ^53^.

## DISCUSSION

In this study, we leveraged large-scale single-cell multi-omics data from CIMA cohort ^27^ to investigate how different BMI categories shape immune cell states and mediate susceptibility to immune-related diseases. By integrating scRNA-seq, scATAC-seq, GRNs, cell type-level xQTLs, and GWAS, we provide a cell type-resolved framework linking BMI-associated molecular alterations to downstream disease risk. Our findings extend prior population-level genetic studies ^4,10–17^ by offering mechanistic insights into how BMI influences immune-related diseases through coordinated regulation of chromatin accessibility, TF activity, and gene expression.

One central observation is the strong cell type specificity of BMI-associated molecular changes. Notably, a substantial proportion of DEGs and DARs were restricted to individual cell types, highlighting notable heterogeneity in immune responses to BMI variation. Mature-NK-dim-FCGR3A, cMono-CD14, gdT2-GZMH, and CD4-Tn-CCR7 subsets consistently exhibited prominent transcriptional and epigenomic perturbations. Together, these findings indicate that regulatory programs in specific immune cell types contribute importantly to BMI-related immune dysregulation.

Functionally, overweight status was associated with a pro-inflammatory immune profile, including enhanced interferon signaling and TNF-related pathways, consistent with the concept of metaflammation ^59–61^. In contrast, underweight status was characterized by distinct immune features, such as altered receptor signaling, antigen recognition, and chemokine-related chromatin accessibility. These findings indicate that underweight and overweight are associated with distinct immune features, which may have different implications for disease risk.

By integrating cis-regulatory inference from scGLUE ^38^ with SMR ^39^-based caQTL–eQTL analyses, we identified numerous peak–gene regulatory links, most of which were cell type specific, while some shared patterns were also observed, such as BMI-associated regulation of *CXCR4* in NK and NKT cells. Additional BMI-associated regulation of genes including *IL32* and HLA family members further highlighted pathways related to inflammation and antigen presentation. To further contextualize these transcriptional changes, we leveraged the GRNs provided by the CIMA resource. Mapping BMI-associated DEGs and DARs onto CIMA GRNs by enrichment analysis revealed 97 TFs, whereas part of them also exhibited differences in motif deviation scores across BMI categories, providing supporting evidence for altered transcriptional activity. Together, the integration of peak–gene links, GRNs, and motif deviation analyses connects BMI-associated TFs, peaks, and downstream gene expression, offering a multi-layered framework to interpret immune variation across BMI categories.

A major strength of this study lies in linking single-cell xQTLs with GWAS of BMI-mediated immune-related diseases. SMR analysis prioritized cMono-CD14, Mature-NK-dim-FCGR3A, and CD8-Tn-CCR7 as key disease-relevant cell types, with RA, asthma, and psoriasis emerging as the most strongly associated diseases. Using RA as a case study, we demonstrated a plausible molecular mechanism underlying increased RA risk in underweight individuals. In MAIT-SLC4A10 cells, increased chromatin accessibility at a *CCR6*-associated regulatory region, together with elevated transcriptional activity of RORC and RORA, was associated with higher *CCR6* expression in the underweight category. Given the established role of *CCR6* in RA pathogenesis, this regulatory axis provides a mechanistic link between low BMI and heightened disease susceptibility.

Several limitations should be noted. Our focus on young adults (20–29 years) minimized age-related confounding but may limit generalizability. In addition, the low prevalence of obesity in the cohort restricted exploration of severe obesity-associated immune alterations.

In summary, our study provides a comprehensive single-cell multi-omics view of how BMI categories modulate immune regulatory landscapes and contribute to immune-related disease risk. By integrating human genetics with cell type-resolved epigenomic and transcriptomic data, we highlight BMI as an active biological modifier of immune function and demonstrate the utility of multi-omics approaches for dissecting complex disease mechanisms.

## METHODS

### Acquisition and Processing of CIMA Cohort Data

We obtained the scRNA-seq and scATAC-seq data of the CIMA cohort in h5ad format from the Trueblood database (https://db.cngb.org/trueblood/cima/), including cell-level expression/accessibility matrices and associated metadata. Using Scanpy ^62^ (version 1.10.4), we subset the data to include 210 samples aged 20–29 years (Supplementary Table 1) for downstream analyses. The SCENIC+ ^57^ based GRN tables of the CIMA cohort, as well as the lists of eGenes and caPeaks for each L4 cell type, were also obtained from the TrueBlood database (https://db.cngb.org/trueblood/cima/). Full summary statistics of CIMA xQTLs used for the SMR analysis were obtained from the OMIX database (accession number: OMIX009786).

### Differential expression analysis

The raw count matrix was normalized using ‘sc.pp.normalize_total’ (with target_sum=1e4) and log-transformed with ‘sc.pp.log1p’ using Scanpy ^62^ (version 1.10.4). To find DEGs between different categories of BMI in different cell types, we applied ‘sc.tl.rank_genes_groups’ based on the non-parametric Wilcoxon rank-sum test. P-value adjustment was performed using benjamini-hochberg correction based on the total number of genes. Genes were considered differentially expressed with Bonferroni corrected p-value <0.05, and an average absolute log₂fold change at least 0.2.

### Differential accessible regions analysis

To find DARs between different categories of BMI in different cell types, the Seurat ^63^ (version 5.3.0) function ‘FindMarkers’ was used. The peak accessibility matrix was normalized by word frequency inverse document frequency (TF-IDF) normalization. As recommended in the Signac ^64^ documentation, logistic regression was employed to test for significance, with total peak counts included as a covariate. P-value adjustment was performed using Bonferroni correction based on the total number of accessible regions. Regions with an adjusted p-value less than 0.05 and average absolute log₂fold change magnitude greater than 0.2 were considered significantly differentially accessible.

### Gene ontology pathway analysis

GO pathways (FDR < 0.05) were identified for DEGs in each cluster using the ‘enrichGO’ function in clusterProfiler ^65^ (version 4.14.6) with org.Hs.eg.db (version 3.20.0). Only GO terms with a fold enrichment greater than 2 were retained. GO pathways (Binom_Adjp_BH < 0.05) were identified for DARs in each cluster using rGREAT ^66^ (version 2.8.0). A background of whole genome hg38 was applied and only GO terms with a fold enrichment greater than 2 were retained.

### Accessible regions annotation

DARs were annotated using the R package ChIPSeeker ^67^ (version 1.42). Annotation of epigenomic datasets using the TxDb database which is from R package TxDb.Hsapiens.UCSC.hg38.knownGene (version 3.22.0).

### scPAFA analysis based on scRNA-seq data

We applied scPAFA ^33^ to quantify pathway activity scores at the single-cell level and to integrate BMI-related differential pathway signals across multiple immune cell types. The pathway sets used as input for scPAFA ^33^ were obtained from Spectra ^68^, which provides a general resource of 231 immunological cell type related and cellular process related gene sets applicable to the analysis of immune-related datasets. In this study, we focused on the 148 broadly applicable cellular process pathways shared across multiple cell types.Cell-level pathway activation scores were computed using the ‘scPAFA.tl.fast_ucell_score’ function based on the UCell ^69^ algorithm. Pseudobulk profiles were then generated using the ‘scPAFA.pb.generate_scpafa_input’ function, with experimental batch and sex included as covariates for adjustment. Subsequently, a Multi-Omics Factor Analysis (MOFA) ^70^ model was trained using the ‘scPAFA.pb.run_mofapy2’ function, with the number of latent factors set to 10.Among the 9 factors inferred by the MOFA model, Kruskal–Wallis tests were performed to identify factors associated with BMI categories, and factor 4 was identified as significantly related to BMI. Pairwise comparisons between BMI categories were further conducted using the Mann–Whitney U test. For factor 4, we used the ‘scPAFA.pl.plot_weights_butterfly’ function to visualize and interpret the high-weight cell type–pathway pairs contributing to this BMI-associated factor.

### Cis-regulatory inference with GLUE feature embeddings

To identify significant cis-regulatory regions for each gene, we employed scGLUE ^38^ (version 0.4.0) for the integration and inference process. First, we identified the top 4000 highly variable genes (HVGs) using ‘pp.highly_variable_genes’ in Scanpy ^62^, and also selected 200,000 features from peak accessibility matrix using the ‘snap.select_features’ function in SnapATAC2 ^71^ (version 2.7.1). Then, gene location information in scRNA-seq data was obtained from the GRCh38.gtf file using the ‘scglue.data.get_gene_annotation’ function. Subsequently, we constructed a prior regulatory graph using the ‘scglue.genomics.rna_anchored_guidance_graph(rna,atac)’ function and mapped HVGs information onto the scATAC-seq data. Dimensionality reduction of the peak matrix from the QC-filtered scATAC-seq data was performed using the LSI algorithm with the ‘scglue.data.lsi’ function and the following parameters: n_components=80, use_highly_variable=True. The raw peak count matrix and LSI dimension matrix from scATAC-seq were used as the ATAC omics input for the GLUE model, configured with the ‘scglue.models.configure_dataset’ function with parameters: prob_model=’NB’, use_highly_variable=True, use_rep=’X_lsi’, and use_batch=’sample’. The scRNA-seq data was downsampled by 50% to reduce data volume and approximately align the number of cells with scATAC-seq. The original gene count matrix and PCA dimension matrix were used as the RNA omics input for the model, configured with the ‘scglue.models.configure_dataset’ function using the parameters: prob_model=’NB’, use_highly_variable=True, use_layer=’counts’, use_rep=’X_pca’, and use_batch=’sample’.

To construct the model, we extracted the highly variable parts of both omics from the prior regulatory graph using the ‘guidance.subgraph’ function to generate guidance_hvf. Model training was performed using the data from both omics along with guidance_hvf by utilizing the ‘scglue.models.fit_SCGLUE’ function with the following parameters: init_kws={’h_dim’:512, ‘random_seed’:666} and fit_kws={’data_batch_size’:2048}. To obtain feature embeddings, we can use the ‘scglue.encode_graph’ method. Given that the GLUE model was trained on highly-variable features, regulatory inference will also be limited to these features. So, we extract the list of highly-variable features for future convenience. The ‘qval’ is obtained by FDR correction of the P-values, and the significant regulatory connections can be extracted based on edge attribute (Q-value < 0.05).

### TF enrichment analysis based on CIMA GRN

The GRN of CIMA comprises 404 eRegulons. Among these, 203 were identified as high-confidence activator eRegulons, in which increased TF activity is associated with positive regulatory effects on downstream targets. To assess the regulatory relevance of these activator eRegulons, we performed enrichment analyses based on DEGs and DARs across BMI categories. Specifically, we used the enricher function in clusterProfiler ^65^, treating each TF as an enrichment term and its associated target peaks and genes as the corresponding gene set. This framework allowed us to evaluate whether the targets of each TF were overrepresented among the DEG/DAR-derived features. TFs with an adjusted p value < 0.05 and at least 20 overlapping targets were considered significantly enriched.

### Motif activity analysis with ChromVAR

We computed motif activities for scATAC-seq data using chromVAR ^72^ (version 1.28.0). Following the chromVAR workflow recommendations, we first constructed a RangedSummarizedExperiment object for each cell type using the ‘SummarizedExperiment’ function. Next, we added GC content bias correction using the ‘addGCBias’ function (with genome = BSgenome.Hsapiens.UCSC.hg38). Peaks were filtered using the ‘filterPeaks’ function to retain only those present in at least one cell.

Position frequency matrices (PFMs) for human TF motifs were retrieved from the JASPAR 2020 ^73^ database via the ‘getMatrixSet’ function in TFBSTools ^74^. These PFMs were then used to annotate peaks with the ‘matchMotifs’ function. Motif deviation scores were calculated using the ‘computeDeviations’ function to quantify deviations from expected chromatin accessibility patterns. To identify differentially active motifs between cell types, we applied the ‘FindAllMarkers’ function (with mean.fxn = rowMeans) to compute the average difference in z-scores. Statistical significance was determined using Wilcoxon rank-sum tests, and p-values were adjusted for multiple testing using benjamini-hochberg correction based on the total number of motifs analyzed.

### TF footprinting

TF footprints were calculated using Signac ^64^’s ‘Footprint’ function and and visualized with the ‘PlotFootprint’ function with the subtract normalization method used to account for Tn5 DNA sequence bias. Motifs selected for footprinting were those enriched in DARs and exhibiting differential motif activity as assessed by ChromVAR.

### SMR analysis

The xQTL summary statistics for eGenes and caPeaks with nominal P were extracted and formatted as BESD files. Additionally, East Asian GWAS summary statistics ^51,52^ for 61 BMI-mediated immune-related diseases (Supplementary Table 10) were utilized in the SMR (version 1.3.1) analysis ^39^. We employed SMR to detect two types of pleiotropic associations within each L4 cell type: (i) variant-caPeak-Disease, (ii) variant-eGene-Disease. The default threshold of 5×10^-8^ was used to select the associated xQTLs as instrumental variables for the SMR test. In each cell type, we applied the Bonferroni method to correct *P*_SMR_ in the SMR results and reject associations with a *P*_HEIDI_ <0.01. Associations with a *P*_SMR_ less than 0.05/number of tests and a *P*_HEIDI_ ≥0.01 were considered significant pleiotropic associations. In addition, significant SMR associations with *P*_GWAS_ >1×10^-5^ were excluded.

## Acknowledgements

We would like to thank all our team members.

## Data and Code availability

The CIMA cohort’s single-cell RNA-seq, single-cell ATAC-seq, xQTL and GRN data were obtained from the CIMA hosted on the TrueBlood database (https://db.cngb.org/trueblood/cima/). Full summary statistics of CIMA xQTLs were obtained from the OMIX database (accession number: OMIX009786). All complete analytical and visualization codes were uploaded to the project repository on GitHub (https://github.com/ZhuoliHuang/CIMA_BMI).

## Author contributions

Z.H., Z.X., J.Y., and C.L. conceived the idea; J.Y., C.L., and P.Y. supervised the work; Z.H., and Z.X. analyzed the data and preformed visualization; all authors wrote the manuscript.

---

**Supplemental Figure. 1.**
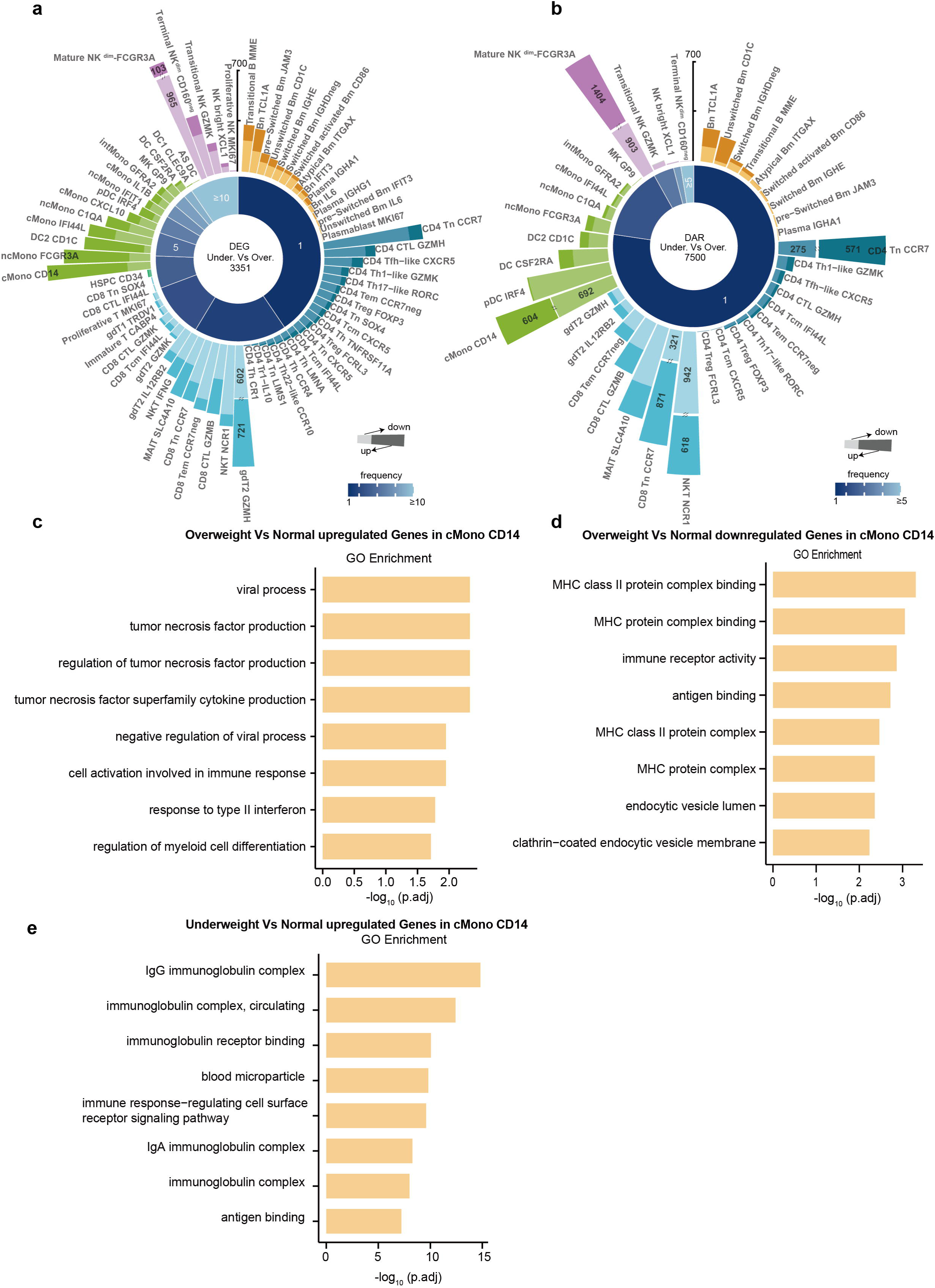
Supplement for BMI-associated DEGs and GO enrichment analysis. (a and b) Bar plot illustrating the number of DEGs detected across 71 cell types (left) and DARs detected across 39 cell types (right) in different BMI categories. The pie chart shows the distribution of significantly detected DEGs or DARs. (c, d and e) GO analysis of DEGs of cMono-CD14 in different comparisons.

**Supplemental Figure. 2.**
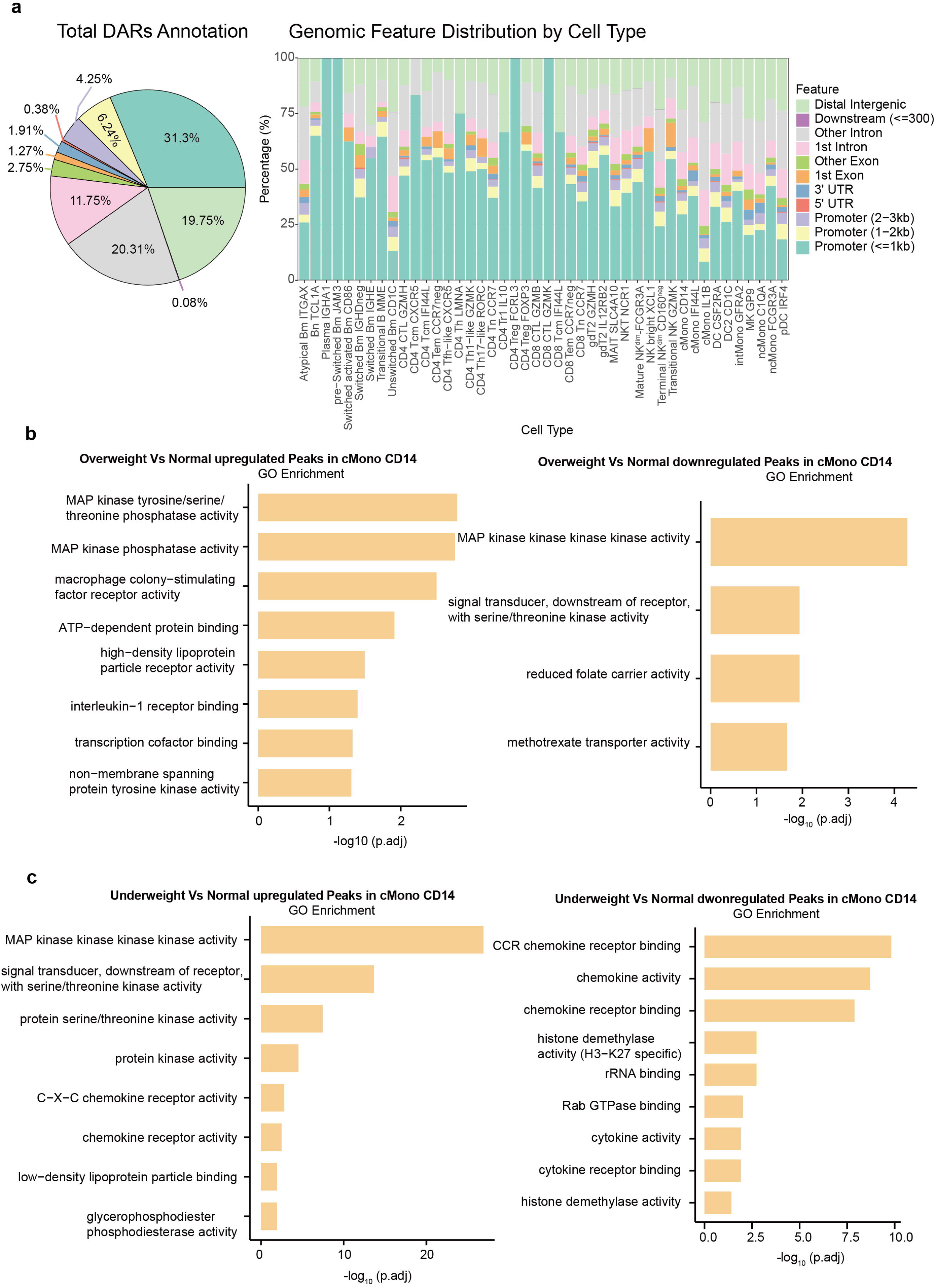
Supplement for BMI-associated DARs and GO enrichment analysis. (a) Pie plot illustrating the distribution of genomic annotations for all DARs (right). Stacked bar plot (left) shows the distribution of DARs based on the peak type in each celltypes. (b and c) GO analysis of DARs of cMono-CD14 in different comparisons.

**Supplemental Figure. 3.**
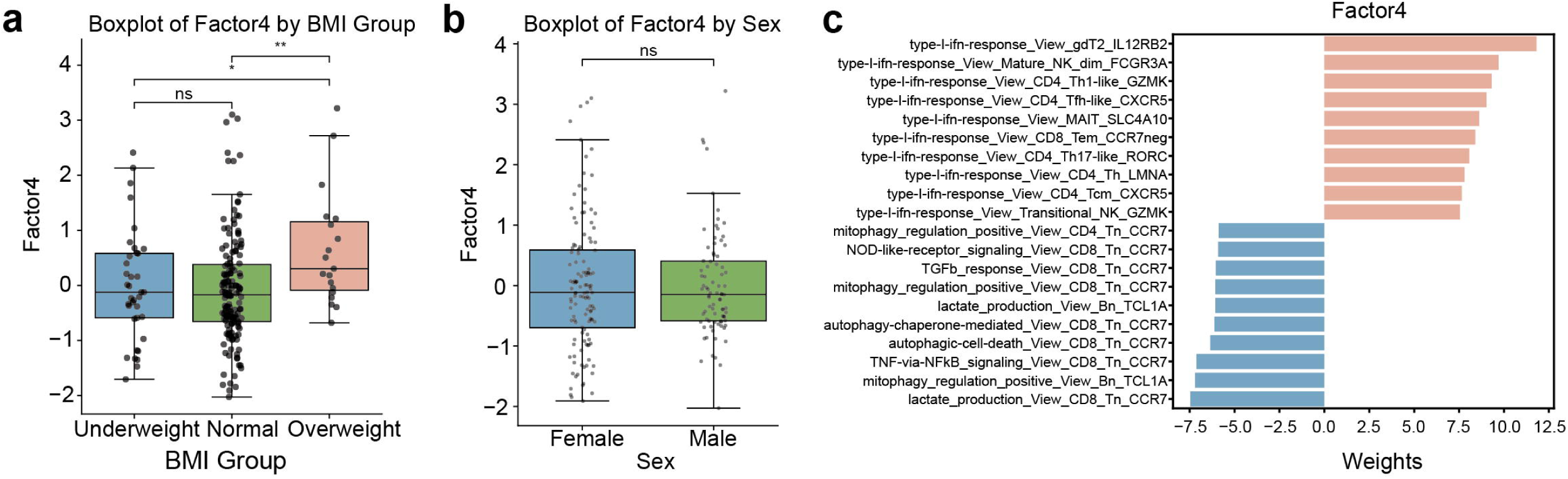
Factor analysis revealed pathways associated with overweight status. (a) Box plots showing the difference in factor 4 values between different BMI categories. (b) Box plots showing the difference in factor4 values between different sex categories. (c) Butterfly bar plot displaying the pathway-cell type pairs with the top 10 positive and negative weights of factors 4.

**Supplemental Figure. 4.**
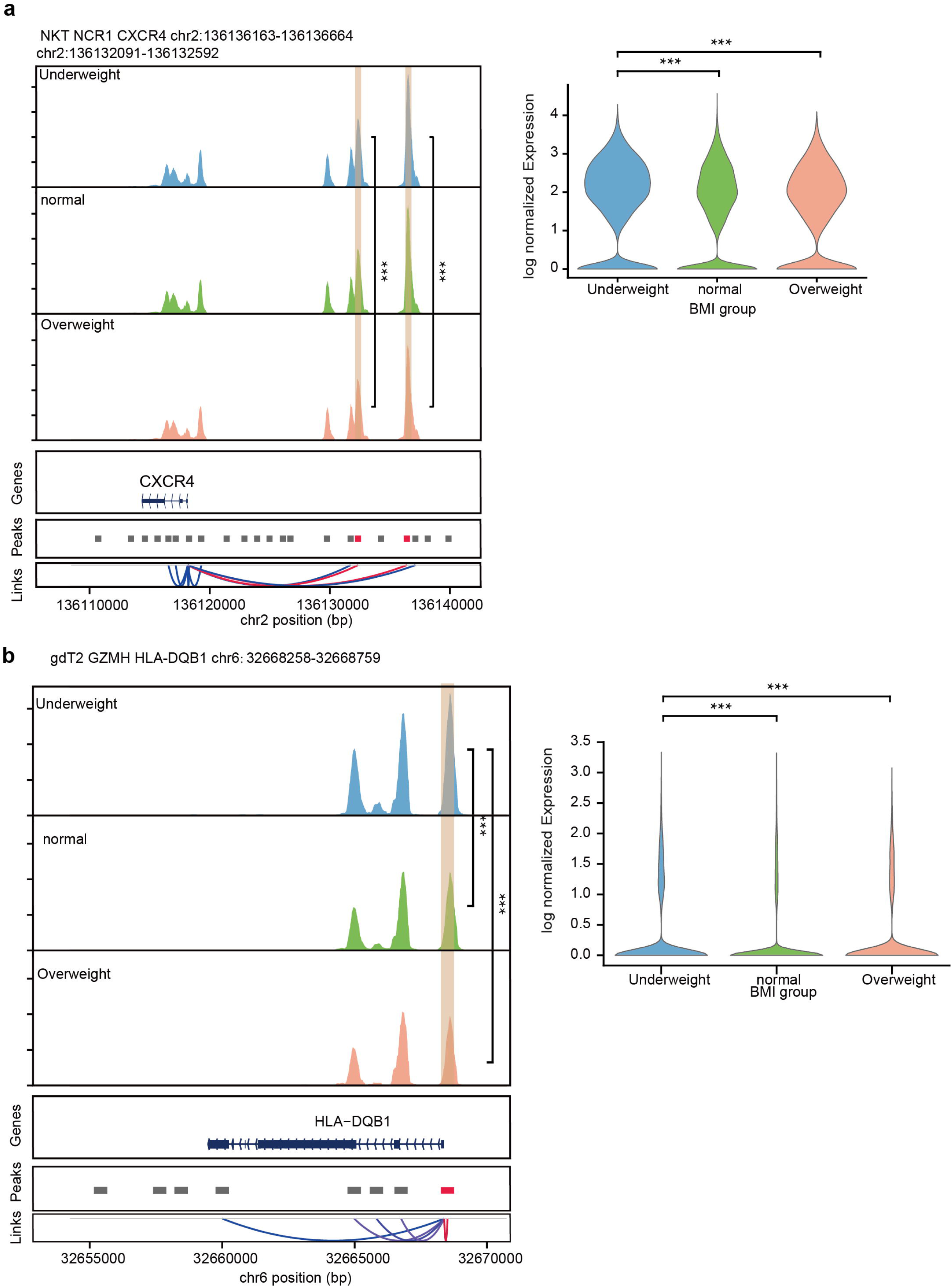
Example of DAR-DEG linkage. (a) Chromatin track of chr2:136136163-136136664 and chr2:136132091-136132592 intersecting CXCR4 gene in different BMI categories in NKT NCR1. The DARs of interest are highlighted in orange. The linkage plot illustrates associations with chromatin accessibility and gene expression which were identified through either SMR-based colocalization or cis-regulatory inference using scGLUE feature embeddings. The link of interest are highlighted in red. Violin plot shows CXCR4 expression in different BMI categories in NKT NCR1. (b) Chromatin track of chr6:32668258-32668759 intersecting *HLA-DQB1* gene in different BMI categories in gdT2-GZMH. The DARs of interest are highlighted in orange. The linkage plot illustrates associations with chromatin accessibility and gene expression which were identified through either SMR-based colocalization or cis-regulatory inference using scGLUE feature embeddings. The link of interest is highlighted in red. Violin plot shows HLA-DQB1 expression in different BMI categories in gdT2 GZMH.

**Supplemental Figure. 5.**
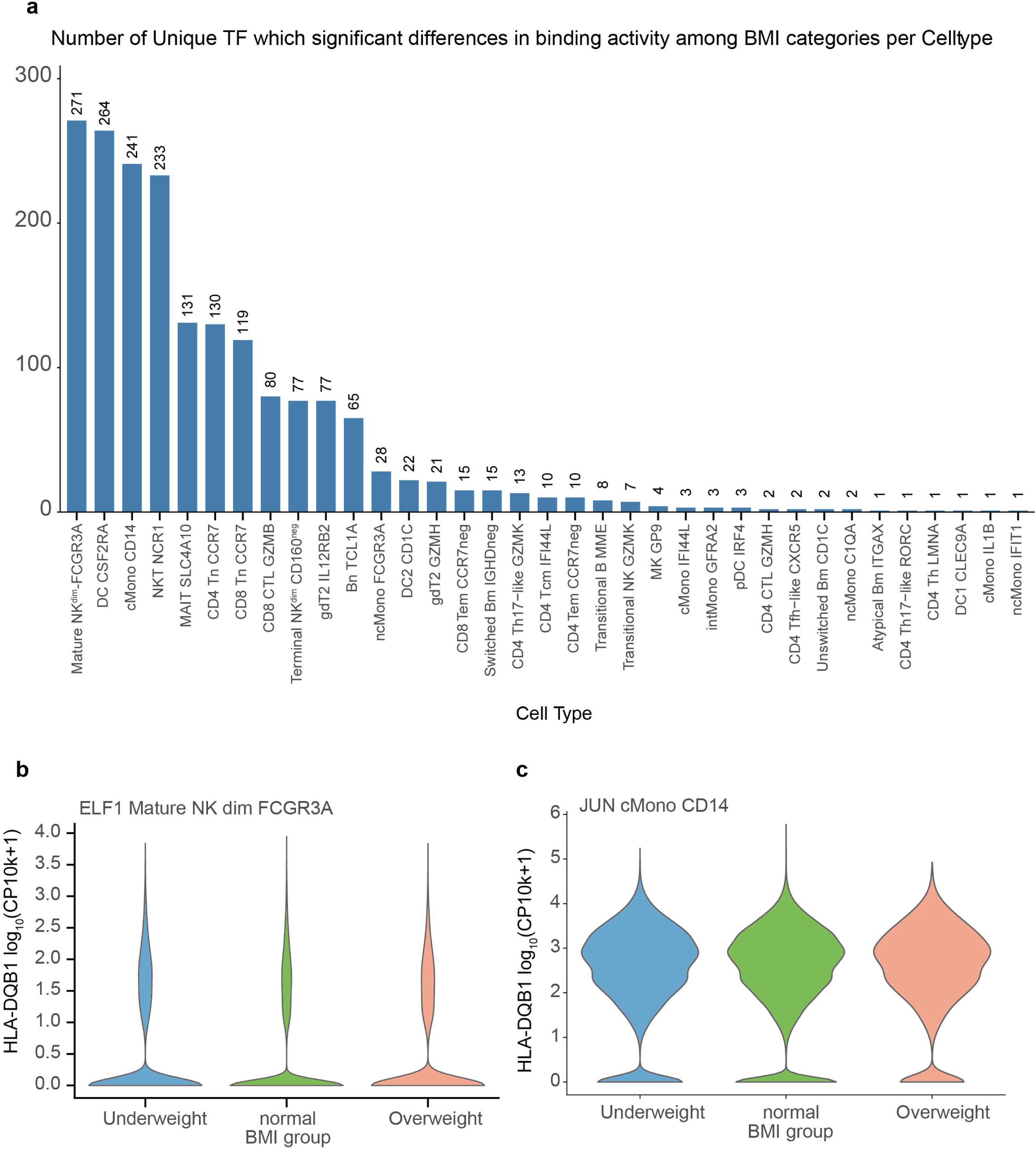
Summary of TF motif deviation results. (a) The bar plot shows the number of TFs with significantly different TF motif deviation z-scores across BMI groups in different celltypes. (b) Violin plot shows *ELF1*expression in different BMI categories in Mature-NKdim-FCGR3A. (C) Violin plot shows *JUN* expression in different BMI categories in cMono-CD14.

**Supplemental Figure. 6.**
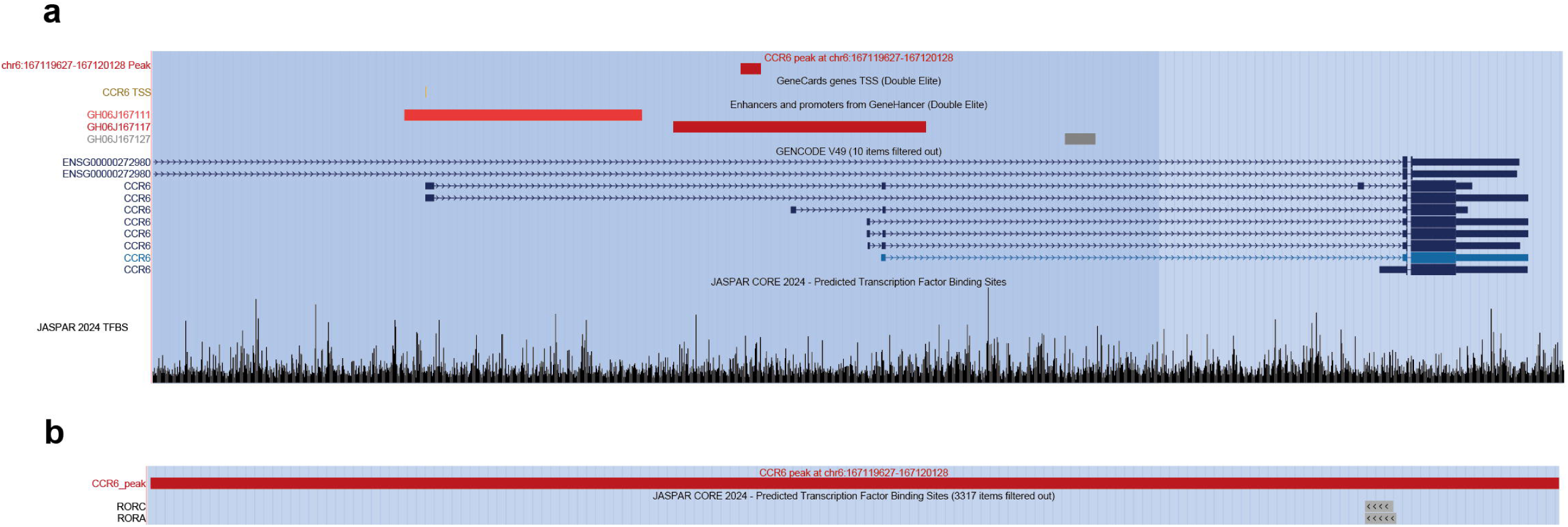
View of *CCR6* region. (a) UCSC Genome Browser (University of California, Santa Cruz, http://genome.ucsc.edu) view of the *CCR6* region showing relevant genetic and regulatory features. (b) UCSC Genome Browser view of chr6:167119627-167120128 peak region.

